## Supplementary Figures And Table for "How accurately can one predict drug binding modes using AlphaFold models?"

### Supplementary Figures and Tables

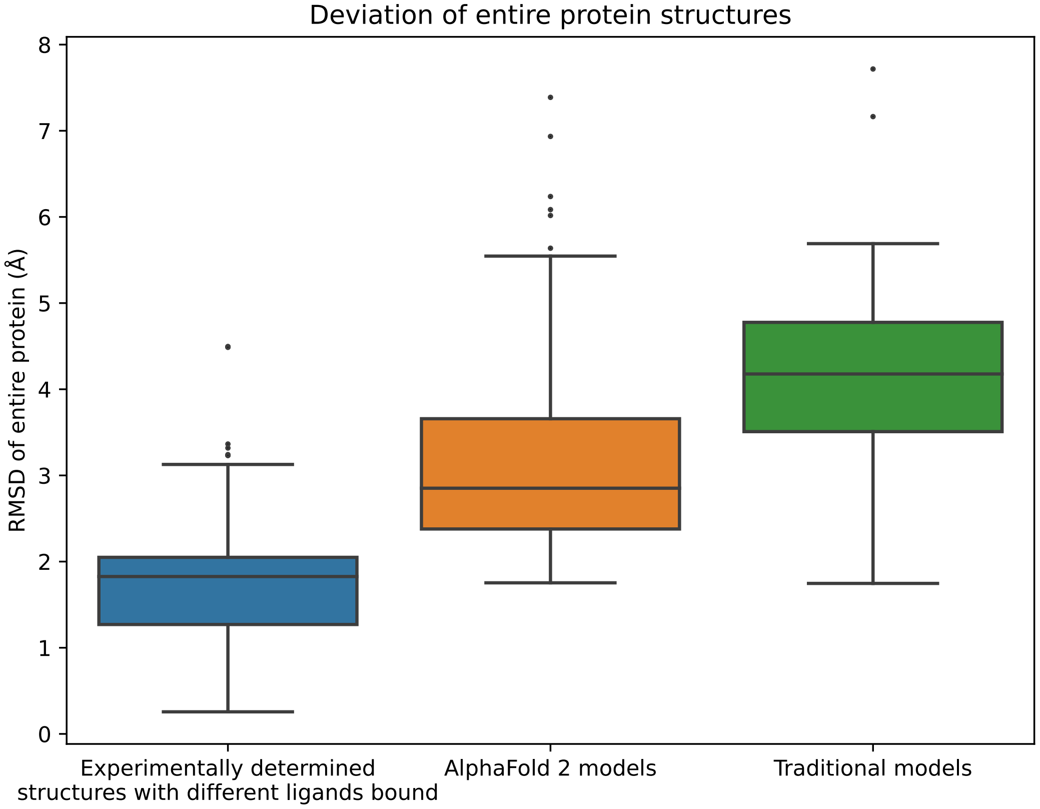


**Figure S1**. *Structural accuracy of modeled proteins.* We compute an all-atom RMSD between each modeled structure and all experimentally determined structures of the same protein. We also compute an all-atom RMSD between each pair of experimentally determined structures of the same protein with different ligands bound. The middle line of each box in the plot is the median RMSD, with the box extending from the 1$^{st}$ to the 3$^{rd}$ quartile and defining the "interquartile range." Whiskers extend to last data points that are within 150% of the interquartile range, and outlier data points beyond those are shown individually.

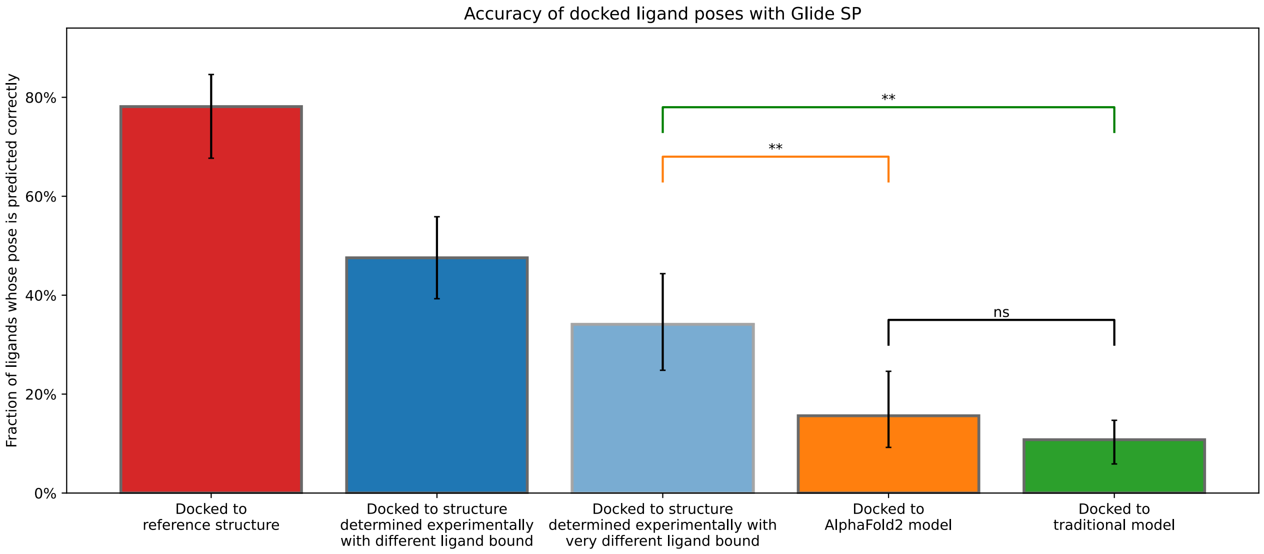


**Figure S2.** *Accuracy of ligand binding poses predicted by computational docking to AlphaFold 2 models, traditional template-based models, or experimentally determined protein structures. This figure is identical to Figure 2 apart from the addition of two bars: (1) The red bar ("docked to reference structure") is for the case where one*docks the ligand from an experimentally determined structure back into the same structure ("self-docking"); (2) the light blue bar is for the case where each ligand is docked only to protein structures determined experimentally in complex with a ligand very different from the one being docked. Very different ligands are defined as ligand pairs in which the atom count of the maximal common substructure that is less than half the atom count of the smaller molecule. Error bars are 90% confidence intervals calculated via bootstrapping. ** for P values < 0.01. ns for P values > 0.05.

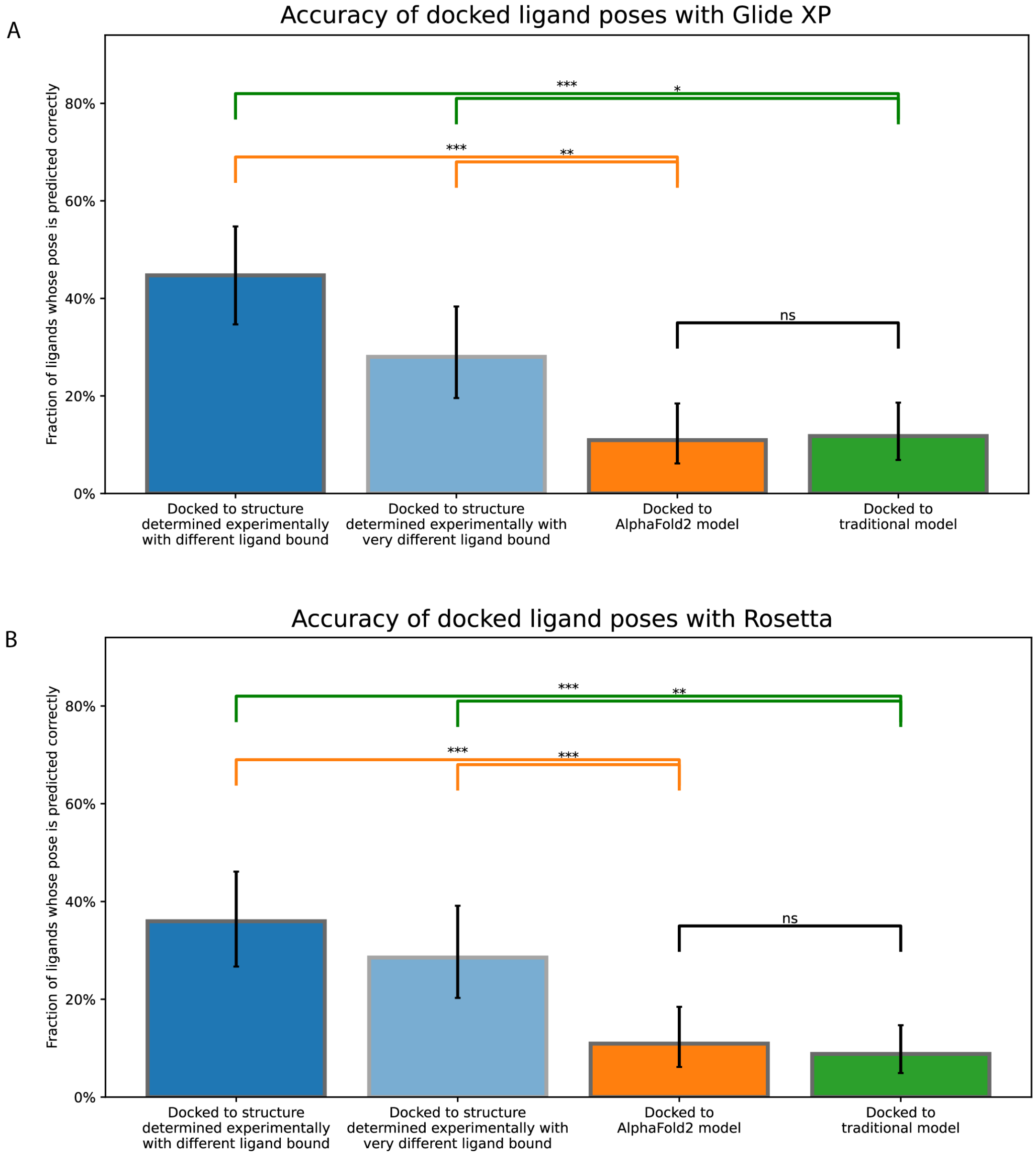


**Figure S3.** *Accuracy of ligand binding poses predicted by computational docking to AlphaFold 2 models, traditional template-based models, or protein structures determined experimentally in complex with ligands different from the one being docked or very different from the one being docked.* These plots are similar to those of Figure 2 and Supplementary Figure S3, except that docking is performance with A) Glide Dock XP and B) Rosetta docking. Error bars are 90% confidence intervals calculated via bootstrapping (see Methods). *** for P values < 0.001, ** for P values < 0.01, * for P values < 0.05. ns for P values > 0.05.

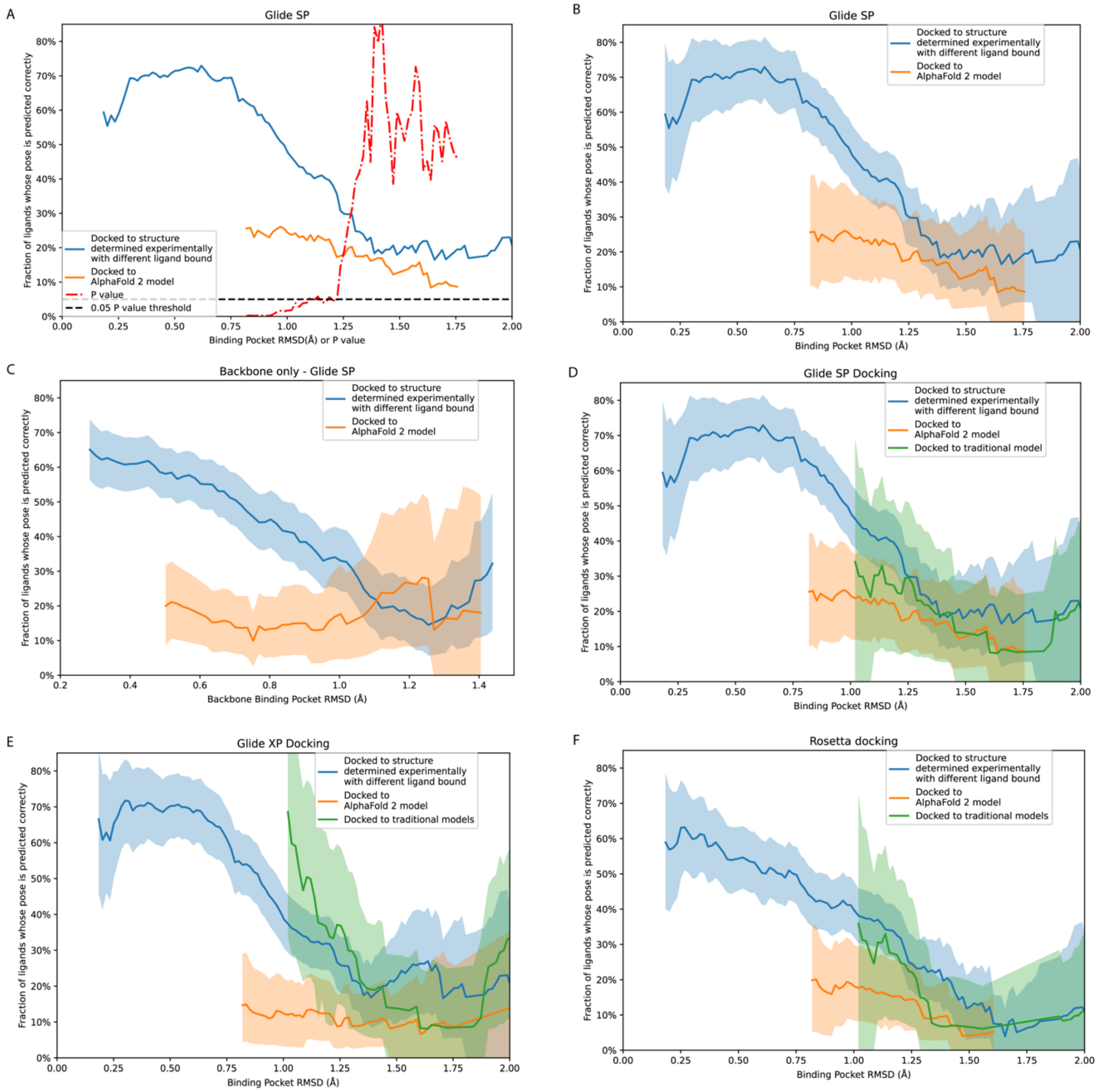


**Figure S4.** *Pose prediction accuracy as a function of binding pocket structural accuracy when docking to AF2 models, experimentally determined structures or traditional template-based models, with different docking protocols*. Using our docking data, we estimate how the pose prediction accuracy of ligand docking varies with the binding pocket all-atom RMSD of the structure being docked into (see Methods). In A) We show calculated P values for the range across binding pocket RMSD. Differences between docking to AF2 and docking to experimentally determined structures are significant for all RMSD values less than 1.1. In B-F) The shaded regions indicate 90% confidence intervals assigned by bootstrapping (see Methods). B) We show the same Glide SP results as in figure 5 but with confidence bounds. In C) we show what this would look like as we examine backbone binding pocket RMSD instead of all atom pocket RMSD. In D-F) we additionally include a curve for docking accuracy with template-based models. D) with Glide SP, E) with Glide XP, and F) with

Rosetta. We consistently see that average pose prediction accuracy using AF2 models is lower than when using experimentally determined structures.

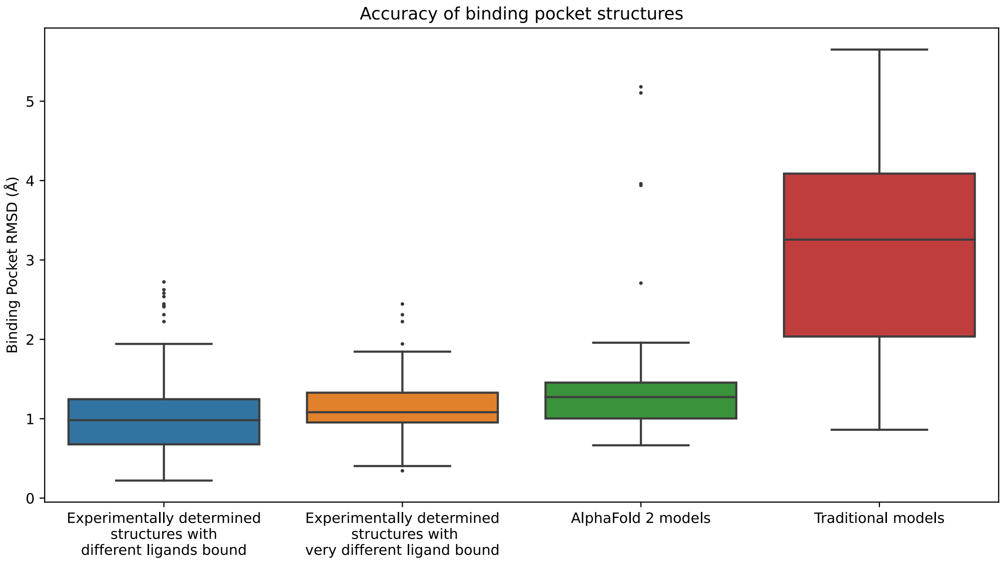


**Figure S5.** *Structural accuracy of modeled binding pockets.* This figure is similar to Figure 1 but includes an additional bar for the case where each ligand is docked only to protein structures determined experimentally in complex with a ligand very different from the one being docked. Very different ligands are defined as ligand pairs in which the atom count of the maximal common substructure that is less than half the atom count of the smaller molecule

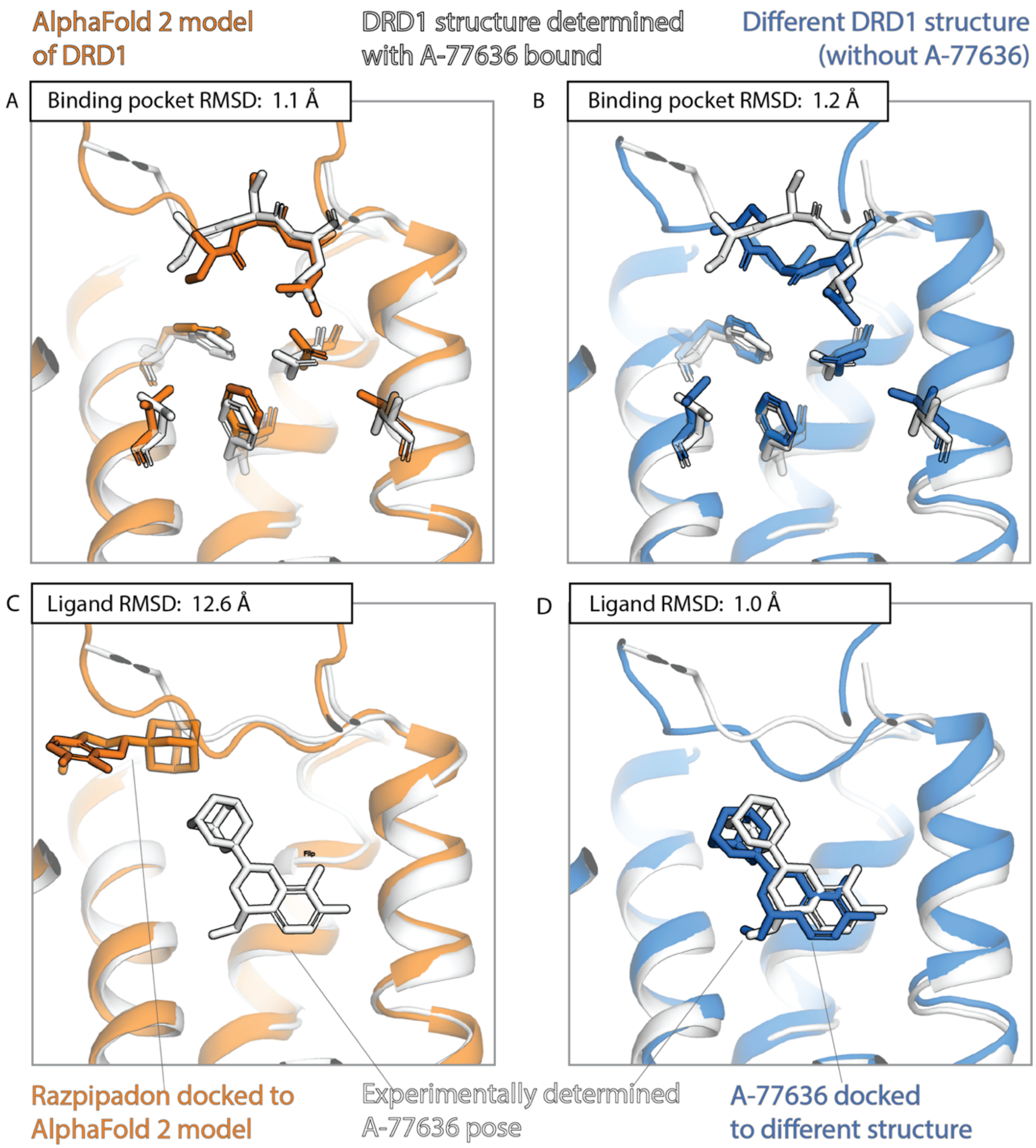


**Figure S6.** An example in which docking to an AF2 model yields poor results even though the model's binding pocket has high structural accuracy. We predict the binding pose of the drug-like ligand A-77636 to the D1 dopamine receptor (DRD1) given either the AF2 model (orange) of DRD1 or the experimentally determined structure (blue) of DRD1 bound to a different ligand, Razpipadon. (A, B) The binding pocket of the AF2 model is more similar (lower RMSD) than the binding pocket of the Razpipadon-bound structure to the binding pocket of the A-77636-bound structure (the "reference structure," white). Amino acid residues whose positions differ most from the reference structure are shown in sticks. (C, D) The A-77636 binding pose predicted by docking is much less accurate (higher RMSD) when using the AF2 model than when using the Razpipadon-bound structure.

*Supplementary Table 1. Structures in benchmark*

| Uniprot ID | Protein name | Class | PDB structure codes |
|---|---|---|---|
| AGRG3 | Adhesion G Protein-Coupled Receptor G3 | B | 7D76,7D77 |
| NK1R | Neurokinin-1 Receptor | A | 6E59,6HLL,6HLO,6HLP,6J20,6J21 |
| 5HT2A | 5-Hydroxytryptamine Receptor 2A | A | 6A93,6A94,6WGT,6WH4,6WHA |
| ADA2A | Alpha-2A adrenergic receptor | A | 6KUX,6KUY |
| CNR2 | Cannabinoid Receptor 2 | A | 5ZTY,6KPC,6KPF,6PT0 |
| DRD1 | Dopamine Receptor D1 | A | 7CKW,7CKX,7CKY,7CKZ,7CRH, 7JOZ,7JV5,7JVP,7JVQ,7LJC,7LJD |
| CLTR2 | Cysteinyl Leukotriene Receptor 2 | A | 6RZ6,6RZ7,6RZ8,6RZ9 |
| CLTR1 | Cysteinyl Leukotriene Receptor 1 | A | 6RZ4,6RZ5 |
| MTR1A | Melatonin Receptor 1A | A | 6ME2,6ME3,6ME4,6ME5,6PS8 |
| MTR1B | Melatonin Receptor 1B | A | 6ME6,6ME7,6ME8,6ME9 |
| PTAFR | Platelet Activating Factor Receptor | A | 5ZKP,5ZKQ |
| TA2R | Thromboxane A2 Receptor | A | 6IIU,6IIV |
| PE2R3 | Prostaglandin E2 Receptor EP2 Subtype | A | 6AK3,6M9T |
| PE2R4 | Prostaglandin E2 Receptor EP4 Subtype | A | 5YHL,5YWY,7D7M |
| PD2R2 | Prostaglandin D2 Receptor 2 | A | 6D26, 6D27 |
| 5HT1A | 5-Hydroxytryptamine Receptor 1A | A | 7E2Y,7E2Z |
| GPBAR | G protein-coupled bile acid receptor 1 | A | 7CFM,7CFN |
| PE2R2 | Prostaglandin E2 Receptor EP2 Subtype | A | 7CX2,7CX3,7CX4 |
